# Real time monitoring of transtibial elevated vacuum prostheses: a case series on socket air pressure

**DOI:** 10.1101/388736

**Authors:** Katherine R. Schoepp, Jonathon S. Schofield, David Home, Michael R. Dawson, Edmond Lou, McNiel Keri, Paul D. Marasco, Jacqueline S. Hebert

## Abstract

Elevated vacuum is a prosthetic suspension method used to reduce slippage between the prosthetic socket and the residual limb. Evaluation of the effectiveness of these systems is limited due to a lack of correlation to actual socket air pressure, particularly during unconstrained movements. This may explain some of the variability in functional outcomes reported in the literature. We developed a light-weight portable socket measurement system to quantify internal socket air pressure, temperature, and acceleration. We implemented the system onto the sockets of three transtibial prosthesis users with mechanical elevated vacuum pumps. Participants completed five functional tasks with and without the vacuum pumps actively connected, including the 2-Minute Walk test, 5-Times Sit-to-Stand test, 4-Square Step test, L-Test, and Figure-8 test. Results demonstrated that the use of elevated vacuum pumps produced different gait profiles and pressure ranges for each user, with significant differences between pump conditions. Two of the participants demonstrated substantially lower air pressure (higher vacuum) over time while the pump was active compared to inactive. The minimum air pressure measured at the completion of the 2-Minute Walk test was −34.6 ± 7.7 kPa, which is not as low as pressures reported in literature during benchtop experiments. One participant did not show substantial changes in pressure over time for either pump condition. Functional task performance was not significantly different between pump conditions. Correlation with accelerometer readings allowed air pressure data to be aligned with the gait cycle; peak positive pressures occurred just following initial contact of the foot in early stance, and the most negative pressures (vacuum) were observed throughout swing. This study has demonstrated the use of a portable data logging tool that may serve the clinical and research communities to quantify the operation of elevated vacuum systems, and better understand the variability of mechanical pump operation and overall system performance.

## Introduction

In 2005, there were an estimated 1.6 million people living with an amputation in the United States; this number is expected to increase to 3.6 million by 2050 (1). Despite advances in prosthetic limb development, optimal socket fit remains a challenge (2–4). Poor suspension may result in slippage between the socket and the residual limb, particularly during the cyclical loading and unloading associated with gait, which can compromise stability (2). This can promote irritation, discomfort, and tissue damage (5). One approach to minimizing this slippage is using elevated vacuum suction suspension, where sub-atmospheric pressure (vacuum) is employed to reduce the relative movement of the user’s residual limb with their prosthetic socket (6). In a typical elevated vacuum socket, the residual limb is covered by a gel liner which sits within a rigid prosthetic socket, and a vacuum is applied through a one-way valve to the space between these layers to improve their connection. The connection between the liner and residual limb is maintained using a proximal seal, which is typically either a suspension sleeve or inner sealing gasket (6). Elevated vacuum systems are predominantly used for attaching lower-limb sockets, though recently there has been preliminary work showing promise for use in transradial (7) and partial-foot amputation cases (8).

Several studies have demonstrated benefits in using elevated vacuum in lower-limb prostheses. When compared to passive suction sockets, vacuum pumps have been shown to maintain or increase residual limb volume during gait (9,10). This may be due to changes in socket-limb interface pressure, where the vacuum reduces positive contact pressures during stance and increases negative air pressures during swing, thereby increasing the fluid drawn into the limb (11). In support of this theory, bioimpedance analysis demonstrated an increase in extracellular fluid volume when walking using a transtibial prosthesis with elevated vacuum (12). Residual limb movement relative to the socket (i.e. pistoning) has been shown to be lower when using elevated vacuum compared to traditional suction and pin-locking systems, with increasing vacuum pressures correlated to reduced pistoning (10,13–15). Improved balance and gait when using elevated vacuum systems has also been demonstrated (10,16,17). Compared to pin-locking and traditional suction sockets, elevated vacuum has demonstrated improved perfusion and preservation of skin barrier function after 16 weeks of use (18). In fact, several studies have found that elevated vacuum systems do not preclude wound healing, and allow patients to ambulate sooner and for longer periods of time compared to other systems (19–21). Generally, elevated vacuum systems are viewed favourably by clinicians, however questionnaire results have shown that they are perceived as being “more expensive, heavier, less durable, and require more maintenance” than a standard socket (22). Several review articles have been published in this area, and while existing evidence for elevated vacuum systems is promising, these reviews have indicated a need for more controlled studies, larger sample sizes, and evaluation of long-term effects (6,23–25).

Recent findings have shown that the level of vacuum (i.e. negative air pressure) is directly related to the amount of pistoning (13), and that changes in pressure may be related to quality of socket fit (26). However, many studies regarding the effectiveness of elevated vacuum do not monitor socket air pressure. Bench-top testing of both electrical pump systems (27,28) and mechanical elevated vacuum systems (27) highlight model-specific differences in measures of performance such as maximum gauge pressure and air evacuation time (27,28). These differences may help to explain variability in study findings, such as in the case series by Sanders *et al.* that found inconsistent results across different elevated vacuum systems (12). Monitoring vacuum pressures while wearing a prosthesis with elevated vacuum could possibly shed light on these differences. For in-lab testing, a pressure monitor (model 2L760, DigiVac, Matawan, NJ) has been used to quantify socket air pressure (27–29). Because this system is tethered to a computer system and comes with the cost of increased bulk and weight it may not appropriate for tasks that require free movement, limiting its use to standing, sitting, and treadmill walking. Xu et al (2017) developed a pressure measurement system to induce a specific vacuum level in order to study the effect on gait parameters, but did not report the changes in vacuum pressure throughout the trials (30). The LimbLogic VS Communicator (Ohio Willow Wood) has been developed to measure socket air pressure in real-time (13,26,31), however it is only designed to interface with the LimbLogic VS system, limiting its usability across a wider range of systems. A discrete monitor that could be used across elevated vacuum systems to measure and log socket air pressure in real-time across of variety of functional tasks could provide valuable quantitative comparisons. To address these limitations, we developed a light-weight portable socket measurement system capable of capturing internal socket air pressure, temperature, and acceleration. This system can either log data to onboard memory, or stream wirelessly and in real-time to a computer. The objective of this paper is to describe the system design, fabrication, and integration of the device, as well as present preliminary results from implementation with three transtibial prosthesis users with mechanical elevated vacuum pumps.

## Methods

Ethics approval was obtained through the University of Alberta’s institutional review board and participants gave written informed consent prior to participation. Three participants currently using an elevated vacuum system in their prosthesis were recruited through prosthetics shops, with details listed in Table 1. Each prosthesis was evaluated by a certified prosthetist and was deemed to be well-fitting at the time of testing. To confirm quality of fit, participants completed the OPUS Lower Extremity Functional Status Measures survey (32), with results ranging from 50 to 70 out of a total possible score of 80.

**Table 1.** Participant information and prosthetic components.

|  | Participant 1 | Participant 2 | Participant 3 |
| --- | --- | --- | --- |
| Participant information |  |  |  |
| Sex | Male | Male | Male |
| Weight | 225 lbs | 175 lbs | 210 lbs |
| Height | 5'11" | 5'10" | 6'2" |
| Amputation information |  |  |  |
| Amputation level | Transtibial | Transtibial | Transtibial |
| Side of amputation | Right | Left | Right |
| Type of amputation | Trauma | Vascular | Vascular |
| Time since amputation | 4 years, 1 month | 3 years, 4 months | 8 years, 4 months |
| Limb length*, geometry | Short, Conical | Medium, Cylindrical | Medium, Cylindrical |
| Wear and performance |  |  |  |
| Hours per day prosthesis worn | 16 | 16 | 16 |
| Days per week prosthesis worn | 7 | 7 | 7 |
| OPUS Functional Status Score | 50 out of 80 | 70 out of 80 | 69 out of 80 |
| Prosthetic components |  |  |  |
| Elevated vacuum system | Harmony P3 (Ottobock, 4R147=K) | Unity Sleeveless Vacuum (Össur) | Triton H (Ottobock, 1C62, Rt. Cat. 3-4-P4N) |
| Liner | Alpha Design Custom Liner (WillowWood, ALC-DES-EO) | Alpha Classic Liner (WillowWood, ALC-5064-E) | Anatomic 3D PUR Liner (Ottobock, 6Y512=265x125-F) |
| Foot | LP Vari-Flex (Össur, 27R C7) | Pro-Flex LP Torsion (Össur, PLTO425L) | Triton H (Ottobock, 1C62, Rt. Cat. 3-4-P4N) |
| Sleeve | Extreme Sleeve (Alps, SFK-28-3) | ProFlex Plus Sleeve (Ottobock, 453A=1-0) | ProFlex Plus Sleeve (Ottobock, 453A40=2-7) |

| Socket type | Thermoplastic temporary socket | Laminated | Environmentally Managed System, Laminated (EMS) |
| --- | --- | --- | --- |

\* Note that “short” limb length refers to a tibia length less than 12 cm and “medium” is between 12 and 15 cm (33)

### Design and Installation of Data Logger

The socket data logger was developed in-house and contained three sensors; an air-pressure sensor (MPXx6250A, Freescale Semiconductor, range: 20 to 250 kPa absolute, reported accuracy: ± 0.25 kPa), an external temperature probe (LMT86, Texas Instruments, range: −50 to 150°C, reported accuracy: ± 0.4°C), and an inertial measurement unit (MPU-9250, InvenSense, range: −8 to 8 G, reported accuracy: ± 0.05 G). Overall dimensions of the device including housing were 18 x 38 x 51 mm with a total weight of 27 g. The device was powered by a lithium polymer battery and communicated using Bluetooth LE in real-time via a custom graphical user interface (GUI) at a frequency of 25 Hz while, simultaneously logging the data to internal on-board memory at the same rate for redundancy.

Air pressure measurements in the socket were obtained by connecting the sensor to the socket via the existing tubing (Participants 1 and 3) or exhaust port connector (Participant 2) between the pump and socket, shown in (Fig 1). A narrow tubing diameter was selected (1/16 inches inner diameter) to ensure that the inclusion of the data logger would have minimal impact on the overall volume of the prosthetic socket; the volume increase for a 20 cm length of tubing is 0.4 cm^3^, relatively small compared to estimated socket volumes ranging from 33 to 197 cm^3^ (28). Therefore, consistent with Boyle’s law and previous prosthetic literature (31), air pressure measurements within this additional tubing are equivalent to the air pressure throughout the prosthetic socket. The temperature probe was placed on the outside of the socket and covered with the prosthetic liner. The housing containing the inertial measurement unit was mounted to the outside of the rigid socket, as was the temperature probe.

**Fig 1.**
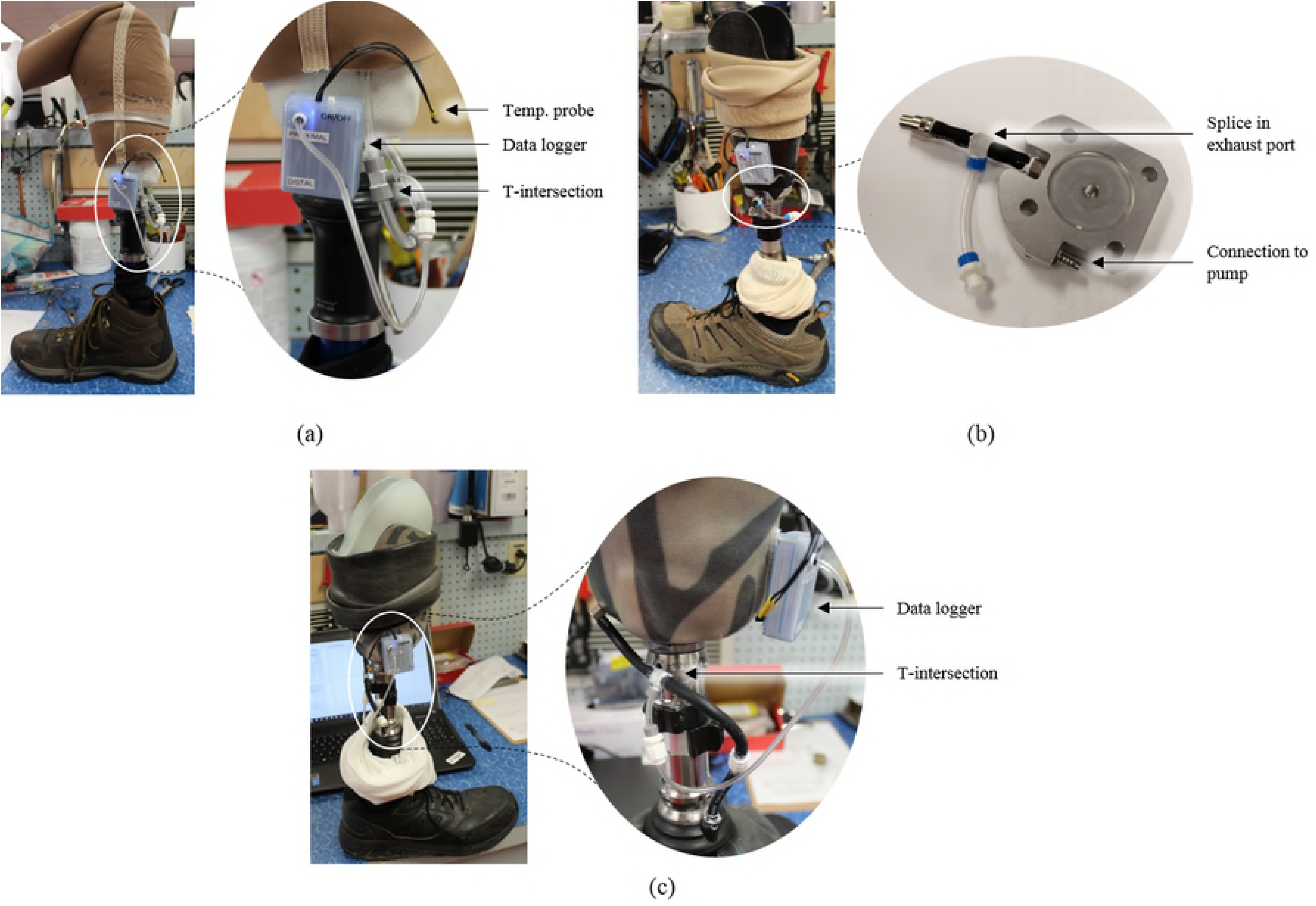
Installation of data logger onto prosthetic socket. Modified sockets for (a) Participant 1, (b) Participant 2, and (c) Participant 3

Pump performance was evaluated in two conditions; active and inactive. In the active condition, the pump was connected to the socket as per manufacturer’s instructions. In the inactive condition, the connection to the pump was replaced by a plug, thereby separating the pump from the socket. Trials were double-blinded; both the participant and researchers did not know the condition of the pump. To ensure the double-blind condition was maintained, a certified prosthetist was responsible for connecting or disconnecting the pump between trials and did not communicate the state of the system until after data analysis was complete. A shroud was placed over the entire prosthetic leg to hide any visual clues. There were however minor differences in auditory cues in the different pump conditions.

### Functional Tasks and Subjective Surveys

Five mobility tasks were performed for each trial in the same order. The first task was a 2-Minute Walk Test, similar to (34), where the participant walked in a large circular hallway (circumference of 190 m) for two minutes. The total distance travelled was measured using a measuring wheel (Rolatape Measuring Systems, Model MM-45M). The participant then completed the Five Times Sit-to-Stand test (35), the 4-Square Step test (36), the L-Test (37), and the Figure-8 test (38); time to task completion, number of steps, and errors were determined from analysis of video footage. At the beginning of the test session, task instructions were provided to the participant and they were given the opportunity to practice each task until comfortable with their performance. This was done to minimize potential learning effects during the trials. During each trial, the 2-Minute Walk test and 5-Times Sit-to-Stand test were completed once, and the 4-Square Step test, L-Test, and Figure-8 test were completed twice.

At the beginning of each trial, the prosthetist connected or disconnected the pump in a room separate from the participant and researchers. Each condition (pump active or inactive) was evaluated twice for a total of four trials, with order of condition block randomized in pairs. Participants were asked to don their prosthesis as usual, then they completed the functional tasks outlined in the order above. If a mistake was made during one of the functional tasks, that specific task was repeated immediately. Once the functional testing was complete, participants were asked to doff their prosthesis, and a seated break of at least five minutes was enforced prior to the next trial.

After each trial the participant completed a short survey to capture their impressions of the prosthesis under the current condition. The survey was modified from the OPUS Satisfaction with Device Score (39), and used a 5-point Likert scale, from 1 (strongly disagree) to 5 (strongly agree), shown in the supplementary file (S1 Table).

### Data Treatment

Analysis was conducted using Excel (Microsoft, 2016) and Matlab (Mathworks, R2017b). Gauge pressure data and acceleration magnitudes were analyzed. For each of the activities, data was broken into individual movements of the gait cycle (i.e. strides) by delineating at maximum pressures within the approximate stride period specified; note that data segments were shifted forward by 10 timesteps (0.4 s) to visually capture the data surrounding the peaks in pressure. The exception is for the sit-to-stand motion where minimum pressures were used to separate movements. The data was then normalized over movement length, creating a scale of 0 to 100% movement completion that allowed for the data to be plotted by condition. Pressure change over time was calculated by determining the slope of the data using simple linear regression. From each individual movement, average and standard deviation values were determined. Two-sample t-tests were used to evaluate differences between active compared to inactive conditions, with variance conditions confirmed using f-tests, and α = 0.05. Note that Trial 4 from Participant 3 was excluded from the analysis due to technical challenges resulting from the donning process that resulted in inadequate seal from the sleeve, compromising the suction suspension.

## Results

Gauge pressure data collected during the 2-Minute Walk test for each participant are shown in Fig 2. The thin lines indicate the real-time pressure measurements, which fluctuated substantially with each stride. The thick lines indicate the overall change in pressure. For Participant 1 (Fig 2a), the pressure remained fairly consistent, regardless of pump condition. For Participant 2 (Fig 2b), in both conditions the pressure dropped (vacuum increased) initially, then stabilized to different values depending on pump condition. For Participant 3 (Fig 2c), the pressure was fairly consistent when the pump was inactive and fell continuously when the pump was active.

**Fig 2.**
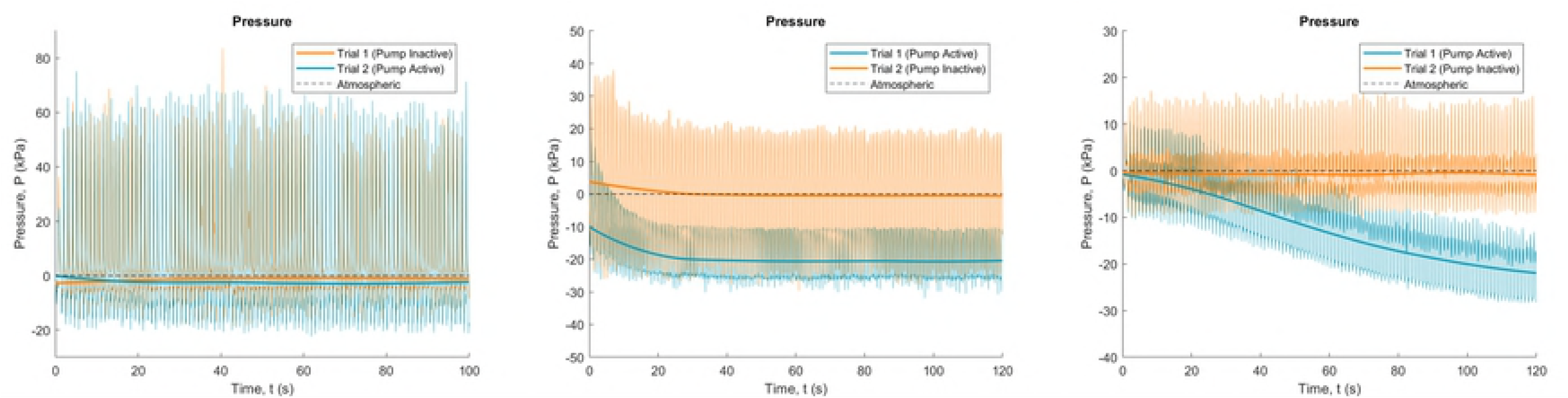
Overall gauge pressure data during 2-minute walk test. Data is shown across (a) Participant 1, (b) Participant 2, and (c) Participant 3. Thin lines indicate raw data and thick lines indicate measurements smoothed using the rloess function in Matlab.

Data for the 2-minute walk test was broken into individual strides as shown in Fig 3. Early strides are indicated in blue with later strides in yellow. This visualization demonstrates differences in vacuum pressures over time. In the case of Participant 1 (Fig 3a), the use of elevated vacuum (pump active) appeared to reduce the variations in pressure occurring with each stride, but not the overall pressures. For Participant 2 (Fig 3b) there was a reduction in overall gauge pressure (increase in vacuum) with subsequent strides in both conditions, with more consistently negative vacuum pressures with the pump active. For Participant 3 (Fig 3c), there was a clear effect of the active pump condition showing progressive reduction in pressures with subsequent strides, compared to the no pump condition which show little to no change.

**Fig 3.**
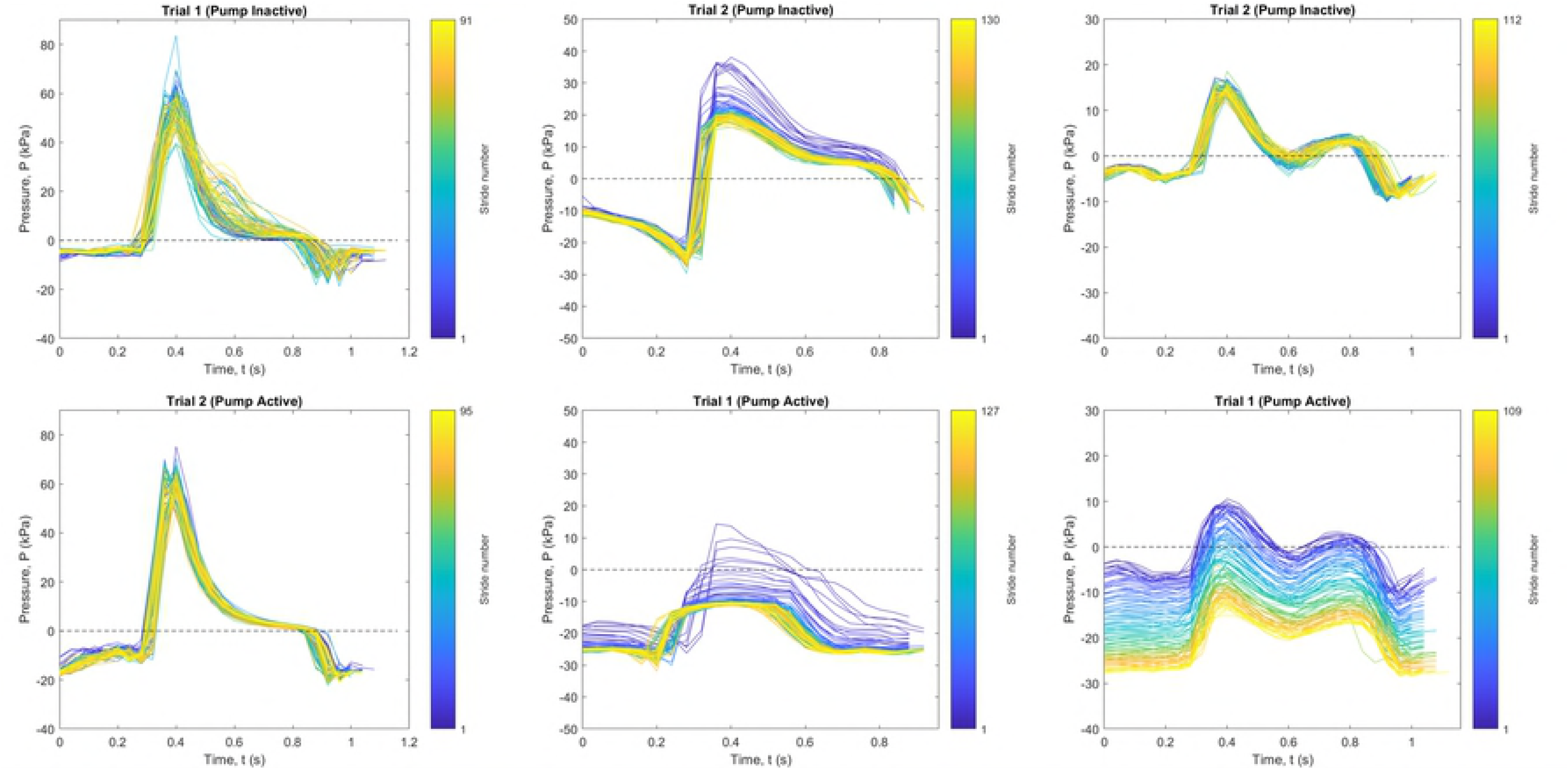
Individual gauge pressure stride data during 2-Minute Walk test. Data is shown with pump inactive (top) and pump active (bottom), across (a) Participant 1, (b) Participant 2, and (c) Participant 3. Blue indicates the first stride and yellow the last, with the legend indicating total stride count.

Average gauge pressure and acceleration data for the 2-Minute Walk test are shown in Fig 4, normalized over the full stride length. As above, Participant 1 (Fig 4a) visually showed small differences in absolute pressure between conditions, and lower pressure variation with the pump active. Participants 2 (Fig 4b) and 3 (Fig 4c) showed large differences in pressure between conditions, where the active elevated vacuum reduced overall pressure. Across all three participants, peaks in pressure were followed by drops; these drops coincided with stable acceleration measurements, where acceleration magnitude was close to 1.0 G.

**Fig 4.**
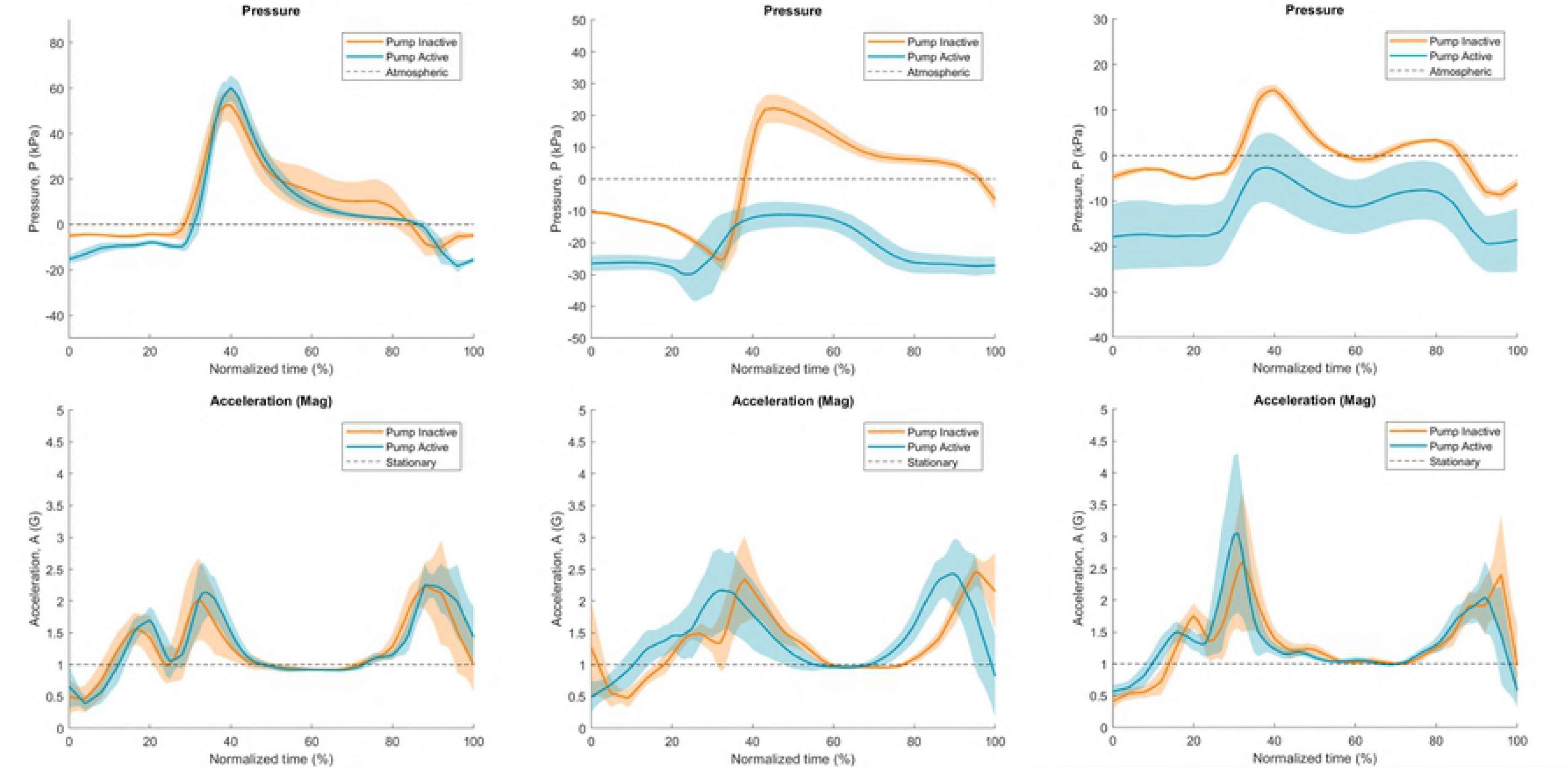
Normalized individual stride data during 2-Minute Walk test. Data is shown for gauge pressure (top) and magnitude of acceleration (bottom), across (a) Participant 1, (b) Participant 2, and (c) Participant 3. Dark lines indicate average measurements, with shaded areas indicating standard deviation.

Values and statistical results from the data analysis of gauge pressure and acceleration for all five tasks are presented in Table 2. There were some significant differences in acceleration magnitude between conditions, though effect size may not be clinically significant. In general, gauge pressure was consistently lower (vacuum higher) while the elevated vacuum system was active, though the effect size varied by participant and activity. Differences between pump active and inactive were smaller during the shorter duration tasks compared to the 2-Minute Walk test.

**Table 2.** Average measured results over individual strides. Reported as mean ± standard deviation. Significant differences are highlighted. Abbreviations are as follows: press. (pressure), accel. (acceleration), mag. (magnitude).

|  |  | Participant 1 |  |  | Participant 2 |  |  | Participant 3 |  |  |
| --- | --- | --- | --- | --- | --- | --- | --- | --- | --- | --- |
|  |  | Inactive | Active | P-value | Inactive | Active | P-value | Inactive | Active | P-value |
| 2-Minute Walk | Gauge press. (kPa) | 8.5 ± 2.9 | 4.6 ± 1.3 | < 0.001 | -0.1 ± 1.4 | -21.7 ± 3.2 | < 0.001 | 0.1 ± 0.3 | -12.4 ± 6.7 | < 0.001 |
| | Accel. mag. (G) | 1.2 ± 0.1 | 1.3 ± 0.0 | < 0.001 | 1.3 ± 0.1 | 1.4 ± 0.1 | < 0.001 | 1.3 ± 0.1 | 1.3 ± 0.1 | 0.743 |
| | Press. change (kPa / min.) | 1.6 ± 0.3 | -0.5 ± 0.2 | 0.016 | -1.6 ± 0.0 | -2.6 ± 0.5 | 0.096 | 0.1 ± 0.2 | -11.7 ± 0.0 | 0.014 |
| 5 Times Sit-to-Stand | Gauge press. (kPa) | 2.7 ± 0.7 | 0.6 ± 0.4 | < 0.001 | -0.7 ± 0.5 | -4.0 ± 2.9 | 0.042 | -1.8 ± 1.1 | -10.1 ± 1.0 | < 0.001 |
| | Accel. mag. (G) | 1.0 ± 0.00 | 1.0 ± 0.01 | 0.193 | 1.0 ± 0.00 | 1.0 ± 0.01 | 0.112 | 1.0 ± 0.01 | 1.0 ± 0.00 | 0.303 |
| 4-Square Step Test | Gauge press. (kPa) | 5.7 ± 4.0 | 6.6 ± 2.9 | 0.264 | 1.3 ± 5.0 | -5.7 ± 4.7 | < 0.001 | 0.8 ± 1.0 | -4.5 ± 1.2 | < 0.001 |
| | Accel. mag. (G) | 1.2 ± 0.13 | 1.1 ± 0.03 | < 0.001 | 1.2 ± 0.12 | 1.1 ± 0.11 | 0.311 | 1.1 ± 0.05 | 1.1 ± 0.04 | 0.919 |
| L-Test | Gauge press. (kPa) | 9.3 ± 3.3 | 7.0 ± 2.9 | < 0.001 | 7.0 ± 6.2 | -11.0 ± 5.6 | < 0.001 | 0.9 ± 0.7 | -4.3 ± 1.1 | < 0.001 |
| | Accel. mag. (G) | 1.3 ± 0.16 | 1.2 ± 0.07 | < 0.001 | 1.3 ± 0.12 | 1.3 ± 0.13 | 0.521 | 1.2 ± 0.08 | 1.2 ± 0.10 | 0.207 |
| Figure-8 Test | Gauge press. (kPa) | 7.7 ± 4.6 | 6.0 ± 1.9 | 0.149 | 3.4 ± 3.4 | -9.3 ± 3.5 | < 0.001 | 0.2 ± 0.6 | -2.9 ± 0.9 | < 0.001 |
| | Accel. mag. (G) | 1.3 ± 0.16 | 1.3 ± 0.12 | 0.433 | 1.2 ± 0.11 | 1.3 ± 0.10 | 0.282 | 1.2 ± 0.06 | 1.2 ± 0.07 | 0.586 |

For further insight into differences with vacuum pressures during the 2-Minute Walk test, we analyzed the first 5 and last 5 strides of the task (Table 3). In all instances, there were significant differences in gauge pressure between the active versus inactive conditions. Differences in pressures between the active and inactive conditions were more pronounced for the last 5 strides of the test compared to the first 5 strides, particularly for Participants 2 and 3. Notably, the maximum air pressure for Participant 2 and 3 for the last 5 strides of the task with the pump active maintained negative values, meaning that the socket was sub-atmospheric throughout the entire gait profile, in contrast to the positive values in the inactive pump condition.

**Table 3.** Average gauge pressure data across initial and final strides during 2-Minute Walk. Reported as mean ± standard deviation. Significant differences are highlighted.

|  |  | Participant 1 |  |  | Participant 2 |  |  | Participant 3 |  |  |
| --- | --- | --- | --- | --- | --- | --- | --- | --- | --- | --- |
|  |  | Inactive | Active | P-value | Inactive | Active | P-value | Inactive | Active | P-value |
| Initial 5 strides | Min. press. (kPa) | -13.8 ± 2.0 | -18.5 ± 1.2 | < 0.001 | -24.9 ± 1.9 | -28.9 ± 5.3 | 0.022 | -9.8 ± 0.7 | -10.1 ± 0.3 | 0.005 |
| | Max. press. (kPa) | 58.1 ± 10.9 | 66.7 ± 7.2 | 0.035 | 38.6 ± 3.6 | 4.3 ± 5.2 | < 0.001 | 15.0 ± 0.9 | 9.6 ± 0.6 | 0.001 |
| Final 5 strides | Min. press. (kPa) | -12.9 ± 1.8 | -20.5 ± 1.7 | < 0.001 | -27.5 ± 3.2 | -34.6 ± 7.7 | 0.002 | -9.3 ± 0.4 | -28.3 ± 0.1 | < 0.001 |
| | Max. press. (kPa) | 54.5 ± 3.4 | 61.8 ± 4.3 | 0.007 | 20.0 ± 1.7 | -12.2 ± 1.8 | < 0.001 | 15.7 ± 0.8 | -13.5 ± 1.0 | < 0.001 |

Temperature measurements of the external socket ranged between 22 and 27°C, however there was no correlation to pump condition.

Functional task performances are summarized in the supplementary material (S2 Table). There were no significant differences in performances based on pump condition (*p* > 0.05 for all comparisons). Functional task performance in both conditions and across all participants fell within normative walking distances during the 2-Minute Walk test (34). Performance during the L-Test and Figure-8 test exceeded reported values based on populations of transtibial amputees and people with mobility disabilities, respectively (37,38). Task durations of the 5 Times Sit-to-Stand and 4-Square Step tests were longer than values reported in normative adult populations (36,40).

Responses to the qualitative survey are summarized in Fig 5. Despite our attempts to blind the participants to the condition of the pump, Participant 1 was able to correctly identify pump condition in all trials. Survey responses of Participant 1 indicated a preference towards the use of the elevated pump, with a perceived improvement in prosthesis fit and comfort, reduced pain and perception of slippage, as well as a greater feeling of control. Participant 2 misidentified the pump conditions for Trials 1 and 2 and was correct for Trials 3 and 4. Participant 3 misidentified every pump condition. Survey responses from Participants 2 and 3 did not indicate a clear preference towards either pump condition.

**Fig 5.**
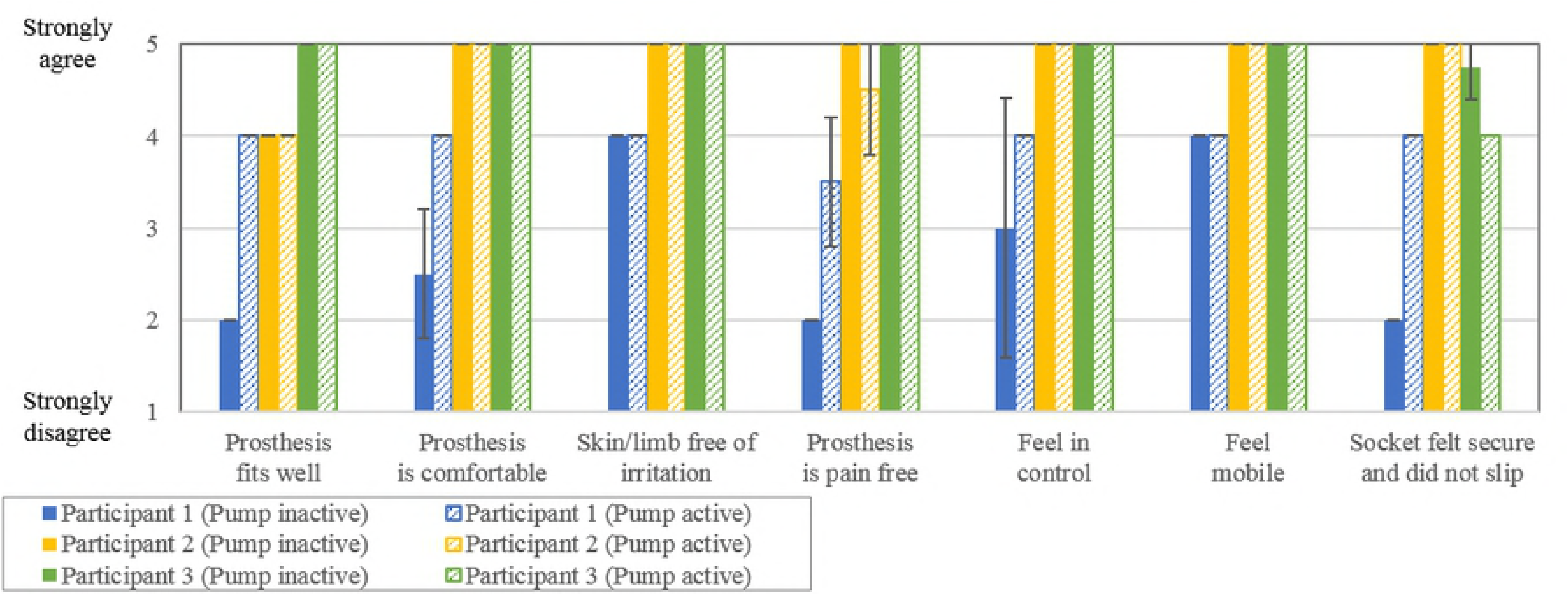
Comparison of qualitative survey scores. Solid colour bars indicate inactive pump condition scores and hatched bars indicate active pump condition scores, with error bars indicating standard deviation

## Discussion

We have developed and applied a device that is able to capture real-time socket pressure and acceleration data while worn non-intrusively and without restricting mobility. This study has demonstrated the performance of the data logger and allowed for the evaluation of air pressure within three different mechanical elevated vacuum systems while worn and performing standardized functional mobility tasks. The use of the pumps resulted in significant changes to socket air pressure over time, where each participant demonstrated different gait profiles and pressure ranges.

With the pump active, both Participants 2 and 3 demonstrated a substantial decrease in socket air pressure over the duration of the 2-Minute Walk test. For Participant 2, the air pressure dropped then plateaued after approximately 50 steps. However, for Participant 3, the pressure dropped continually over the duration of the trial. The air pressure in both participants’ sockets reached a similar final negative pressure range at the end of the task. With the pump inactive, the air pressure values at the end of the trials were similar to those at the beginning; the maximum pressure continued to fluctuate above atmospheric pressure. However, with the pump active, air pressure readings were consistently negative for the final 5 strides, indicating that the elevated vacuum was maintaining a sub-atmospheric pressure throughout the full stride. Interestingly, these two participants had difficulties distinguishing when the pump was active, and questionnaire results did not show a clear preference towards either condition. It was surprising that the participants with apparently effective vacuum systems could not accurately detect when the pump was active.

In contrast, the air pressure in Participant 1’s socket did not change substantially over the 2-Minute Walk test, in both active and inactive pump conditions. However, with the pump active there was a more consistent pressure profile. This participant was the most successful at correctly identifying pump condition, and questionnaire results indicated a strong preference towards the pump being active. Differences between participants may be due to a combination of factors other than pump design, including socket fit and material properties, limb geometry, donning process, and gait pattern, to name a few. In particular, this participant had a short conical limb with less soft tissue coverage, in comparison to the other 2 participants. Future work should investigate the effect of soft tissue compliance and volume on effectiveness of vacuum systems.

Similar pressure trends were seen during the other walking-based tasks (Figure-8 and L-test), though they were not as pronounced due to the shorter task duration. Some average acceleration magnitudes were significantly different between conditions; however, these differences were very small and may not be clinically relevant. The 5-Times Sit-to-Stand activity yielded lower variance in pressure and acceleration compared to other activities, likely because the prosthetic leg was planted against the floor rather than suspended from the limb in swing phase.

Using the measured acceleration readings, changes in socket air pressure can be roughly correlated to phases of the gait cycle. Willemsen *et al.* demonstrated that stable accelerations equivalent to gravity correspond to stance, variable readings to swing, and that peaks occur during push off and foot down (41). We can therefore infer that peak positive pressures occurred just following initial contact of the foot in early stance, with pressure decreasing during stance. The most negative pressures (highest vacuum) were observed throughout swing. This is similar to the pressure profiles measured by Chino *et al.*, where nine transtibial amputees using suction sleeves were evaluated (42). It may be valuable in future work to synchronize the pressure profiles with specific phases of the gait cycle.

Pressure ranges reported in literature vary depending on the type of pump used. Our measured results did not attain negative pressures as low as the benchtop testing conducted by Komolafe et al. (28), which evaluated different mechanical pumps using a material testing system and found that a vacuum pressure of −57.6 kPa could be achieved in less than 50 loading cycles or 80 seconds. Xu et al (30) recommended a moderate level of 50 kPa to optimize comfort and gait symmetry; their pressures were manually pulled rather than induced by the mechanical pump. The minimum pressures observed at the completion of the 2-Minute Walk test during our study were −34.6 ± 7.7 kPa. This may be due to differences between idealistic ‘bench-top’ testing conditions and real-world prosthetic sockets worn by participants; loading profiles and rates were substantially different, and it is likely that the seal of a socket on an amputated limb is inferior to an idealized system. Chino *et al.* found minimum pressures using a suction sleeve ranged between −7 and −31 kPa over ten gait cycles (42). In contrast, air pressures created by electrical elevated vacuum pumps have been reported to range between −27 to −85 kPa (9–11,13,19,20,26,27,29,31). This disparity indicates there may be large differences in air pressure between systems, and more evaluation is needed to better understand the impact of these differences on prosthesis user function.

Functional task performances were compared to reported data. All three participants met or exceeded reported scores for the walking-based tasks (2-Minute Walk, L-Test, and Figure-8) but demonstrated reduced performance for both the 5 Times Sit-to-Stand and 4-Square Step tests. There were no significant differences in task performances between pump conditions, suggesting that the short-term use of elevated vacuum may not have a measurable impact on functional mobility. This is in contrast with the more longer term study conducted by Samitier et al. (17) that found improvements in functional task scores of 16 transtibial participants after 4-weeks of training with an elevated vacuum system, when compared to their previous system.

## Future Work

The data logger tool, testing protocols, and analyses presented contribute to the clinical and research communities by helping to quantify the operation of elevated vacuum systems, and to bridge the gap between the measurement of mechanical pump operation and overall system performance and function.

This study quantified socket air-pressure across three elevated vacuum systems within worn prosthetic sockets, during specific tasks allowing unconstrained walking in the clinical lab environment. The next step is to study trends over longer periods of time outside of the laboratory or clinical setting. This will allow further inferences regarding the pressure changes that occur within a socket not only during various movements, but also during daily living tasks and throughout a longer wearing time. The on-board acceleration measurements could be used to determine compliance, activity level, or to monitor falls, as suggested in literature (43,44). The temperature sensor may provide insight into the impact of different environments on vacuum performance; inclusion of a second temperature probe could allow differences in internal socket temperature and environmental temperature to be studied. It may also be interesting to integrate our system with additional sensors, such as limb-socket interface pressure sensors or strain gauge sensors to quantify forces and moments applied to the prosthesis and residual limb (44).

Future work may also involve determining correlations between socket air pressure and measurements of the residual limb. For example, residual limb volume loss has been demonstrated in seated (9) and standing (12) tasks. It would be valuable to understand how this loss may be related to average pressures versus cyclical pressure changes, and also to different soft tissue characteristics of residual limbs. It will be valuable to investigate pressure differences between mechanical and electrical pumps, as electrical pumps may be set to greater vacuum pressures than the mechanical pumps evaluated in this study, and will likely generate different pressure profiles.

In the future, these techniques should be useful for clinicians, developers, and researchers to address questions related to elevated vacuum system performance. As a clinical tool, this could be used to quickly identify leaks in a socket, understand user compliance, and determine trends in performance throughout the day. There is also a potential for benefit in remote health-care applications (44), or to provide information to developers regarding the impact of various prosthetic components and manufacturing techniques.

## Acknowledgements

The authors would like to thank the Glenrose Rehabilitation Hospital for providing space to conduct the testing as well as the following prosthetics shops for their assistance with participant recruitment: Edmonton Prosthetic Services, Northern Alberta Limb, and the Prosthetic and Orthotic Care Company. They also acknowledge Ossur for loaning the plate (Ossur Icelock 544 Plate Unity) used to collect data for Participant 2. Thank you to Stéphane Magnan and Fraaz Kamal for their assistance with developing the data logger.

## Supporting Information

**S1 Table. Qualitative survey questionnaire.** Modified from the OPUS Satisfaction with Device Score (39).

**S2 Table. Functional modbility task results.** Functional task performance across entire trial, reported as mean ± standard deviation.

## References

1 Ziegler-Graham K, MacKenzie EJ, Ephraim PL, Travison TG, Brookmeyer R. Estimating the prevalence of limb loss in the United States: 2005 to 2050. Arch Phys Med Rehabil 2008 Mar;89(3):422–429.

2 Mak AFT, Zhang M, Boone DA. State-of-the-art research in lower-limb prosthetic biomechanics-socket interface: A review. J Rehabil Res Dev 2001;38(2):161–173.

3 Legro MW, Reiber G, del Aguila M, Ajax MJ, Boone DA, Larsen JA, et al. Issues of importance reported by persons with lower limb amputations and prostheses. J Rehabil Res Dev 1999 Jul;36(3):155–163.

4 Klute GK, Kantor C, Darrouzet C, Wild H, Wilkinson S, Iveljic S, et al. Lower-limb amputee needs assessment using multistakeholder focus-group approach. J Rehabil Res Dev 2009;46(3):293–304.

5 Levy W. Chapter 26: Skin Problems of the Amputee. In: Bowrker JH, Michael JW, editors. Atlas of Prosthetics: Surgical, Prosthetic, and Rehabilitation Principles: Mosby Year Book; 1992.

6 Kahle JT, Orriola JJ, Johnston WJ, Highsmith MJ. The effects of vacuum-assisted suspension on residual limb physiology, wound healing, and function: A systematic review. Technology & Innovation 2014;15:333–341.

7 Lang M, Coyers D. Utilization of negative pressure for socket suspension in upper extremity prosthetics. MEC 2008.

8 Arndt B, Caldwell R, Fatone S. Use of a partial foot prosthesis with vacuum-assisted suspension: A case study. J Prosthet Orthot 2011;23(2):82–88.

9 Goswami J, Lynn R, Street G, Harlander M. Walking in a vacuum-assisted socket shifts the stump fluid balance. Prosthet Orthot Int 2003 Aug;27(2):107–113.

10 Board WJ, Street GM, Caspers C. A comparison of trans-tibial amputee suction and vacuum socket conditions. Prosthet Orthot Int 2001;25(3):202–209.

11 Beil TL, Street GM, Covey SJ. Interface pressures during ambulation using suction and vacuum-assisted prosthetic sockets. J Rehabil Res Dev 2002 Nov-Dec;39(6):693–700.

12 Sanders JE, Harrison DS, Myers TR, Allyn KJ. Effects of elevated vacuum on in-socket residual limb fluid volume: Case study results using bioimpedance analysis. J Rehabil Res Dev 2011;48(10):1231–1248.

13 Gerschutz MJ, Hayne ML, Colvin JM, Denune JA. Dynamic effectiveness evaluation of elevated vacuum suspension. J Prosthet Orthot 2015;27(4):161–165.

14 Klute GK, Berge JS, Biggs W, Pongnumkul S, Popovic Z, Curless B. Vacuum-assisted socket suspension compared with pin suspension for lower extremity amputees: effect on fit, activity, and limb volume. Arch Phys Med Rehabil 2011 Oct;92(10):1570–1575.

15 Darter BJ, Sinitski K, Wilken JM. Axial bone-socket displacement for persons with a traumatic transtibial amputation: The effect of elevated vacuum suspension at progressive body-weight loads. Prosthet Orthot Int 2016;40(5):552–557.

16 Kuntze Ferreira AE, Neves EB. A comparison of vacuum and KBM prosthetic fitting for unilateral transtibial amputees using the Gait Profile Score. Gait Posture 2015;41(2):683–687.

17 Samitier CB, Guirao L, Costea M, Camos JM, Pleguezuelos E. The benefits of using a vacuum-assisted socket system to improve balance and gait in elderly transtibial amputees. Prosthet Orthot Int 2016 Feb;40(1):83–88.

18 Rink C, Wernke MM, Powell HM, Gynawali S, Schroeder RM, Kim J, et al. Elevated vacuum suspension preserves residual-limb skin health in people with lower-limb amputation: Randomized clinical trial. J Rehabil Res Dev 2016;53(6):1121–1132.

19 Hoskins RD, Sutton EE, Kinor D, Schaeffer JM, Fatone S. Using vacuum-assisted suspension to manage residual limb wounds in persons with transtibial amputation: A case series. Prosthet Orthot Int 2014;38(1):68–74.

20 Traballesi M, Delussu AS, Fusco A, Iosa M, Averna T, Pellegrini R, et al. Residual limb wounds or ulcers heal in transtibial amputees using an active suction socket system. A randomized controlled study. Eur J Phys Rehabil Med 2012;48(4):613–623.

21 Brunelli S, Averna T, Delusso S, Traballesi M. Vacuum assisted socket system in transtibial amputees: Clinical report. Orthopädie-Technik Quarterly 2009:2.

22 Gholizadeh H, Lemaire ED, Eshraghi A. The evidence-base for elevated vacuum in lower limb prosthetics: Literature review and professional feedback. Clin Biomech 2016/07;37:108–116.

23 Safari MR, Meier MR. Systematic review of effects of current transtibial prosthetic socket designs-Part 1: Qualitative outcomes. J Rehabil Res Dev 2015;52(5):491–508.

24 Safari MR, Meier MR. Systematic review of effects of current transtibial prosthetic socket designs-- Part 2: Quantitative outcomes. J Rehabil Res Dev 2015;52(5):509–526.

25 Gholizadeh H, Lemaire ED, Eshraghi A. The evidence-base for elevated vacuum in lower limb prosthetics: Literature review and professional feedback. Clin Biomech (Bristol, Avon) 2016 Aug;37:108–116.

26 Wernke MM, Schroeder RM, Haynes ML, Nolt LL, Albury AW, Colvin JM. Progress Toward Optimizing Prosthetic Socket Fit and Suspension Using Elevated Vacuum to Promote Residual Limb Health. Adv Wound Care (New Rochelle) 2017 Jul 1;6(7):233–239.

27 Major MJ, Caldwell R, Fatone S. Comparative effectiveness of electric vacuum pumps for creating suspension in transfemoral sockets. J Prosthet Orthot 2015;27(4):149–153.

28 Komolafe O, Wood S, Caldwell R, Hansen A, Fatone S. Methods for characterization of mechanical and electrical prosthetic vacuum pumps. J Rehabil Res Dev 2013;50(8):1069–1078.

29 Major MJ, Caldwell R, Fatone S. Evaluation of a Prototype Hybrid Vacuum Pump to Provide Vacuum-Assisted Suspension for Above-Knee Prostheses. J Med Device 2015 Dec;9(4):0445041–445044.

30 Xu H, Greenland K, Bloswick D, Zhao J, Merryweather A. Vacuum level effects on gait characteristics for unilateral transtibial amputees with elevated vacuum suspension. Clin Biomech (Bristol, Avon) 2017 Mar;43:95–101.

31 Gerschutz MJ, Haynes ML, Colvin JM, Nixon D, Denune JA, Schober G. A vacuum suspension measurement tool for use in prosthetic research and clinical outcomes: Validation and analysis of vacuum pressure in a prosthetic socket. J Prosthet Orthot 2010;22(3):172–176.

32 Heinemann AW, Bode RK, O’Reilly C. Development and measurement properties of the Orthotics and Prosthetics Users’ Survey (OPUS): a comprehensive set of clinical outcome instruments. Prosthet Orthot Int 2003 Dec;27(3):191–206.

33 Arwert HJ, van Doorn-Loogman MH, Koning J, Terburg M, Rol M, Roebroeck ME. Residual-limb quality and functional mobility 1 year after transtibial amputation caused by vascular insufficiency. J Rehabil Res Dev 2007;44(5):717–722.

34 Bohannon RW, Wang YC, Gershon RC. Two-minute walk test performance by adults 18 to 85 years: normative values, reliability, and responsiveness. Arch Phys Med Rehabil 2015 Mar;96(3):472–477.

35 Guralnik JM, Simonsick EM, Ferrucci L, Glynn RJ, Berkman LF, Blazer DG, et al. A Short Physical Performance Battery Assessing Lower Extremity Function: Association With Self-Reported Disability and Prediction of Mortality and Nursing Home Admission. J Gerontol 1994 03/01;49(2):M85–M94.

36 Dite W, Temple VA. A clinical test of stepping and change of direction to identify multiple falling older adults. Arch Phys Med Rehabil 2002 Nov;83(11):1566–1571.

37 Deathe AB, Miller WC. The L test of functional mobility: measurement properties of a modified version of the timed “up & go” test designed for people with lower-limb amputations. Phys Ther 2005 Jul;85(7):626–635.

38 Hess RJ, Brach JS, Piva SR, VanSwearingen JM. Walking skill can be assessed in older adults: validity of the Figure-of-8 Walk Test. Phys Ther 2010 Jan;90(1):89–99.

39 Shirley Ryan AbilityLab. Satisfaction With Device Score. Orthotics Prosthetics Users Survey 2015.

40 Bohannon RW. Reference values for the five-repetition sit-to-stand test: a descriptive meta-analysis of data from elders. Percept Mot Skills 2006 Aug;103(1):215–222.

41 Willemsen AT, Bloemhof F, Boom HB. Automatic stance-swing phase detection from accelerometer data for peroneal nerve stimulation. IEEE Trans Biomed Eng 1990 Dec;37(12):1201–1208.

42 Chino N, Pearson JR, Cockrell JL, Mikishko HA, Koepke GH. Negative pressures during swing phase in below-knee prostheses with rubber sleeve suspension. Arch Phys Med Rehabil 1975 Jan;56(1):22–26.

43 Redfield MT, Cagle JC, Hafner BJ, Sanders JE. Classifying Prosthetic Use via Accelerometry in Persons with Trans-Tibial Amputations. J Rehabil Res Dev 2013;50(9):1201–1212.

44 Hafner BJ, Sanders JE. Considerations for development of sensing and monitoring tools to facilitate treatment and care of persons with lower-limb loss: a review. J Rehabil Res Dev 2014;51(1):1–14.

